# Sporadic, global linkage disequilibrium between unlinked segregating sites

**DOI:** 10.1101/030247

**Authors:** Daniel A. Skelly, Paul M. Magwene, Eric A. Stone

**Affiliations:** Department of Biology, Duke University, Durham, North Carolina 27708, USA; Department of Biological Sciences, North Carolina State University, Raleigh, North Carolina 27695, USA

**Author notes:** Correspondence to: Eric A. Stone.

## Abstract

Demographic, genetic, or stochastic factors can lead to perfect linkage disequilibrium (LD) between alleles at two loci without respect to the extent of their physical distance, a phenomenon that Lawrence *et al*. (2005a) refer to as “genetic indistinguishability”. This phenomenon can complicate genotype-phenotype association testing by hindering the ability to localize causal alleles, but has not been thoroughly explored from a theoretical perspective or using large, dense whole-genome polymorphism datasets. We derive a simple theoretical model of the prevalence of genetic indistinguishability between unlinked loci, and verify its accuracy via simulation. We show that sample size and minor allele frequency are the major determinants of the prevalence of perfect LD between unlinked loci but that demographic factors, such as deviations from random mating, can produce significant effects as well. Finally, we quantify this phenomenon in three model organisms and find thousands of pairs of moderate-frequency (> 5%) genetically indistinguishable variants in relatively large datasets. These results clarify a previously underexplored population genetic phenomenon with important implications for association studies, and define conditions under which it is likely to manifest.

---

A basic genomic property of sexually reproducing organisms is genetic disequilibrium, the non-random association between alleles at two or more loci. This quantity is often referred to as linkage disequilibrium (LD) to emphasize the important role of genetic linkage in generating and maintaining the association at physically proximal loci. The concept of LD as a property of two loci that are physically near each other is a natural viewpoint given the large body of theoretical and empirical work characterizing mechanisms by which demography and selection influence patterns of LD between physically proximal loci. The existence and extent of LD between linked loci is influenced by the demographic history of a population, reveals locus-specific selective forces, and is a critical parameter governing the resolution of marker-trait association studies (reviewed in Slatkin, 2008). Nevertheless, it is well known that physically distant or even independently segregating loci can also exhibit non-random associations. Specifically, LD between alleles segregating at physically unlinked loci can be induced by a variety of forces such as selection, genetic drift, non-random mating, epistasis, pleiotropy, and non-random chromosome transmission (Michie, 1953; Bennett and Binet, 1956; Nei, 1967; Crow and Kimura, 1970; Lewontin, 1988; Petkov *et al*., 2005; Platt *et al*., 2010; Rohlfs *et al*., 2010; Corbett-Detig *et al*., 2013; Long *et al*., 2013). In this paper we examine non-random associations between alleles at pairs of physically unlinked loci. We use the term LD to describe these associations but stress that they are not a result of genetic linkage between the loci.

For two main reasons, a common approach to empirical examinations of LD in genome analysis studies has been to focus on characterizing LD across modestly-sized bins of specified physical distance (e.g. The International HapMap Consortium, 2005; Liti *et al*., 2009). First, this has been an informative approach for exploring the well-recognized tradeoffs that LD engenders in mapping studies. Specifically, surveys of LD within bins allow assessment of the degree to which LD may facilitate the identification of risk-conferring variants through the typing of nearby neutral polymorphisms but may simultaneously impede efforts to finely map the causal variants responsible for those signals (Kruglyak, 1999; Sutter *et al*., 2004). Second, in genome studies cataloging a large number of polymorphic sites, exploration of LD across the complete genome often constitutes a daunting set of calculations, since the number of possible pairs of single nucleotide polymorphisms (SNPs) increases as *n*^2^ where *n* is the number of SNPs.

A number of previous studies have explored LD between unlinked loci. Rohlfs *et al*. (2010) observed association between alleles present at two putatively coevolving human gamete-recognition genes, and found support for the hypothesis of selection for allele matching at these loci. Similarly, Takano-Shimizu *et al*. (2004) reported LD between polymorphisms in *Drosophila* chemoreceptor genes, which they attributed to the effects of multilocus selection with epistasis. Sved (2011) and Koch *et al*. (2013) examined measures of LD between blocks or patches of variants in subsets of the HapMap dataset, with Sved (2011) finding a slight bias toward positive associations among unlinked blocks and Koch *et al*. (2013) observing an excess of LD extending over long ranges on the same chromosome. However, the focus of these authors on measures of LD between blocks of variants necessarily obscures the most extreme examples of LD between specific pairs of unlinked sites. Petkov *et al*. (2005) focused on LD “domains” in mice consisting of large blocks of sequence with disequilibrium between relatively distant markers on the same chromosome, and they identified an example of LD between domains on two different chromosomes containing genes with strong functional similarity, implying selection for coadapted alleles (Petkov *et al*., 2005). The authors also identified additional cases of high interchromosomal LD (*D*′ > 0.8) but did not focus on this result and did not discuss in detail the frequency of high interchromosomal LD or the distribution of LD among loci with strong disequilibrium. Long *et al*. (2013) calculated genome-wide pairwise LD in a dense polymorphism dataset and found many pairs of loci with high *r*^2^ values even after correcting for population structure, but did not focus on modeling this phenomenon or quantifying the factors that could affect it. Lawrence *et al*. (2005a) examined LD between ~30,000 SNPs on chromosome 20 in several human populations, focusing on SNPs in perfect LD that they dubbed “genetically indistinguishable” SNPs (giSNPs; we adopt this abbreviation and use it throughout this paper). The authors considered both linked and unlinked variation, and found that most giSNPs occurred within haplotype blocks, but did not explore the underlying causes leading to the emergence of unlinked giSNPs (Lawrence *et al*., 2005a). Finally, we note that Lawrence *et al*. (2009) published a tool for examining genome-wide LD in the HapMap dataset.

The investigations described herein are motivated by prospects for genomewide association mapping studies (GWAS) in model and non-model organisms. Specifically, falling sequencing costs and methodological advances are making it increasingly feasible to gather virtually complete catalogs of genomic variation in moderate-sized samples of individuals from a species of interest. This raises the possibility of performing GWAS in a setting where the casual allele(s) are almost certainly tested directly. This approach ameliorates the concerns that several previous studies have raised involving tests for indirect association using so-called “tag” SNPs that are in LD with untyped variants (Zhang *et al*., 2004; Terwilliger and Hiekkalinna, 2006). However, the approach becomes potentially problematic when the possibility exists for very strong LD between alleles at a true causal site and at unlinked loci elsewhere in the genome. If alleles at a causal site and at one or more physically distant loci were to form a cluster of SNPs in perfect LD, it would be impossible to localize the causal site using marker-trait association. At worst, a causal site that is contained within such a cluster along with distant loci that happen to be near *a priori* candidate genes could falsely reinforce assumptions about the genetic basis of trait variation.

This potential pitfall for association studies has been noted previously in the literature. In a review article, Lawrence *et al*. (2005b) emphasized the potential difficulties that giSNPs could create for localizing causal variation in the association mapping setup. They gave an unambiguous example of this phenomenon appearing in an early empirical exploration of the potential for gene expression quantitative trait locus (QTL) mapping in humans (although they acknowledged that their sample size was small; Lawrence *et al*., 2005b). Rohlfs *et al*. (2010) raised the possibility of this same phenomenon and called for further investigation of interchromosomal LD, although they focused on the role of selection. The possibility of LD between unlinked loci hindering localization of genes in association mapping was also discussed in a study of LD in cattle, although the magnitude of LD between unlinked loci was lower than what we consider (Farnir *et al*., 2000). In the *Drosophila* literature, several published genome-wide association studies provide empirical examples of unlinked SNPs with low association *p*-values and either perfect or high long-range LD (Jordan *et al*., 2012; Harbison *et al*., 2013; Swarup *et al*., 2013).

This study has several factors that differentiate it from previous work. First, we consider all (or nearly all) polymorphic sites present in large genomic datasets consisting of complete genomes for at least 100 individuals, and we calculate LD between all pairs of sites no matter their physical distance. Second, we focus our attention on the most extreme values of LD between physically distant loci; only these pairs of sites will pose the type of problem that we describe above in the GWAS setting. We specifically examine perfect LD (genetic indistinguishability) as this phenomenon is amenable to simple theoretical modeling and to demonstrate that this “worst-case” phenomenon is in fact present in real data. Finally, we explore how genetic indistinguishability could be caused purely by stochastic or demographic factors, as such factors are frequently unaccounted for in studies of the genetic basis of phenotypic variation.

To address these issues, we first derive a simple theoretical model to quantify the rate of giSNP occurrence in a randomly mating population, and verify its accuracy via coalescent simulations. Next we show that sample size and minor allele frequency (MAF) are the most important factors determining the prevalence of giSNPs, but that demographic forces and deviations from random mating can also affect the rate of giSNP occurrence, in some cases dramatically. Finally, by fully enumerating the global landscape of LD in several large model organism datasets, we demonstrate that a significant fraction of loci in real datasets can be members of giSNP clusters. As complete sequence data for cohorts of individuals becomes readily available in many model and non-model organisms, it will be important to recognize that this phenomenon can have important implications for the success of trait mapping studies.

## Materials and Methods

### Simulations

We performed simulations to validate our theoretical model and to explore the effects of population structure and demographic perturbations on the prevalence of giSNPs using the software `ms` (Hudson, 2002). Since we are primarily interested in LD between unlinked sites, we assume that segregating sites arise from independent realizations of the Wright-Fisher process. Thus, for all simulations with more than one segregating site, we chose a single site at random and ignored the remaining segregating sites. Then, the collection of *N* sites resulting from *N* independent simulations with at least one segregating site represented the hypothetical overall collection of genomic variants. For all simulations we used a sample size of 50 chromosomes (haploid individuals). We used a mutation rate of *μ* = 1 × 10^−10^ per base pair.

### Population bottleneck simulations

In simulations exploring population bottlenecks, the initial effective population size was *N_e_* = 10,000. We simulated a bottleneck that occurred 100 generations before population sampling. After the reduction in population size due to the bottleneck, the population size remained constant (at the new, reduced size). The severity of the bottleneck varied, ranging from a drastic reduction in population size (1% of original size) to no reduction (100% of original size).

### Exponential growth simulations

In simulations of exponential growth, population expansion began 50 generations before population sampling. Starting population size varied, and for all simulations the ending effective population size was *N_e_* = 10, 000. Varying the starting population size allowed us to simulate exponential growth that differed in magnitude. Starting population size ranged from very small (*N_e_* = 50, very strong exponential growth) to large (*N_e_* = 10, 000, no exponential growth).

### Population splitting simulations

For simulations modeling a single population splitting into two subpopulations, we considered a population with effective size *N_e_* = 10,000 split into two subpopulations, each with size *N_e_* = 5, 000. We varied the timing of the population split from 0 to 10,000 generations, but once the split occurred it was absolute and no migration between subpopulations was allowed. For our ending sample of 50 chromosomes we drew *N* = 25 chromosomes from each population.

### Ten population model

We simulated a single population of size *N_e_* = 10,000 that fragmented into ten subpopulations 500 generations before population sampling. Each subpopulation had an equal size, *N_e_* = 1, 000. For any given simulation, migration between each of the ten subpopulations was symmetric and occurred at a constant rate from the fragmentation event until the present. The migration rate varied between simulations. At the lowest level there was no migration allowed between subpopulations, and at the highest level migration replaced 90% of each subpopulation each generation. For our ending sample of 50 chromosomes we drew *N* = 5 chromosomes from each population.

### Calculating linkage disequilibrium

For simulated, theoretical, and empirical data, we use the term allelic configuration to refer to the specific constellation of individuals carrying the minor and major allele at a segregating site. To identify giSNPs, we converted genotype data to counts specifying the number of minor alleles present in each individual, and identified sites with identical allelic configurations. For empirical data, we ignored missing data and identified giSNPs using only pairwise complete data (that is, only individuals with non-missing genotype data for both loci of interest). To measure the entropy of the distribution of allelic configurations, we used the formula − ∑*_i_ p_i_* log_2_ *p_i_* where *i* indexes the allelic configurations observed for a particular MAF and *p_i_* is the fraction of all allelic configurations of that MAF that possess configuration *i*. This is a standard formulation for measuring entropy in an information theoretic context. For a particular MAF, the entropy is independent of sample size (the number of polymorphic sites) given a constant probability of sampling each of the possible allelic configurations. Furthermore, entropy is a more appropriate measure of the evenness of the distribution of allelic configurations than an alternate measure such as the standard deviation, because the former measure does not require assigning arbitrary numerical values to allelic configurations. We performed all analyses using R (R Core Team, 2014) version 3.1.0 and python version 2.7.3 (Python Software Foundation, 2014).

### Genetically indistinguishable SNPs in real data

For our empirical examinations of giSNPs, we focused on biallelic SNPs, and discarded any sites with over 10% missing data. When tabulating cases of perfect LD, we only examined loci that were on different chromosomes or far enough apart on the same chromosome to be effectively randomly assorting. Specifically, we obtained rough estimates of the meiotic recombination rate [in centimorgans (cM) per kilobase (kb) or megabase (Mb)] and converted this to the approximate physical distance at which loci on the same chromosome are 50 cM apart.

For *S. cerevisiae,* we used data from the “Yeast 100 Genomes” collection described in Strope *et al*. (2015). We tabulated LD between SNPs at least 100 kb apart or on different chromosomes, since the meiotic recombination rate in *S. cerevisiae* is approximately 0.5 cM/kb (Cherry *et al*., 1997).

For *D. melanogaster,* we used freeze 2.0 samples from the *Drosophila melanogaster* Genetic Reference Panel project (Huang *et al*., 2014). The meiotic recombination rate in *D. melanogaster* is on the order of 10 cM/Mb (Comeron *et al*., 2012), so we ignored giSNPs less than 5 Mb apart on the same chromosome. The individuals in this project have been inbred to near complete homozygosity, but a few heterozygous sites still persist, so we set genotypes at these sites to missing data and treated all genotypes as haploid.

For *A. thaliana,* we used genotype data (non-imputed data) of 180 lines from Sweden (Long *et al*., 2013). The meiotic recombination rate in *A. thaliana* is roughly 3.6 cM/Mb (Salome *et al*., 2012), so we ignored giSNPs less than 14 Mb apart on the same chromosome.

## Results and Discussion

### A theoretical model for the prevalence of genetically indistinguishable variants

To gain a detailed understanding of the phenomena that affect the prevalence of giSNPs in an idealized haploid population, we derived a simple theoretical model. We assume that samples are drawn from a single panmictic, constant-sized Wright-Fisher population, with mutations generated according to the infinite-sites model, and that all mutations are selectively neutral. Since we are interested in LD between unlinked sites, we assume that segregating sites arise from independent realizations of the Wright-Fisher process.

We begin by considering a variant with MAF *k*/*N*, where *N* is the number of samples drawn from the haploid population. For a set of variants drawn from the above model, the expected fraction of variants with the same MAF *k/N* is approximately

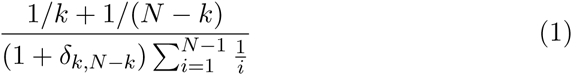

where *δ* is the Kronecker delta. Equation 1 was previously derived by Tajima (1989, eqn. 51) and Fu (1995, eqns. 6-8).

We use the term allelic configuration to refer to the specific constellation of individuals carrying the minor and major allele at a segregating site (i.e. which individuals carry the minor allele and which carry the major allele). Thus, two sites are giSNPs if and only if they share identical allelic configurations. The presence of identical allelic configurations does not imply an identical derived allele distribution among samples, since a derived allele with frequency *k/N* or (*N* − *k*)/*N* could present the same allelic configuration.

Under the neutral Wright-Fisher model (panmixis, constant population size, infinite-sites mutation), for a given MAF *k/N*, all allelic configurations are equally likely since the labeling of individuals is arbitrary. There are 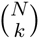 unique allelic configurations for a site with MAF *k/N*. Thus, the fraction of variants with minor allele frequency *k/N* that have an identical allelic configuration is 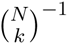.

Combining these results, given a variant with MAF *k/N*, the expected fraction of other variants that are genetically indistinguishable can be expressed as

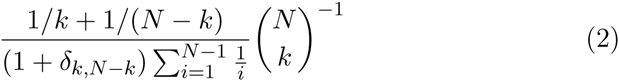

This expression represents the probability that, with a single variant in hand, a second variant drawn at random will be genetically indistinguishable from the first (i.e. they are giSNPs). This probability is independent of the total number of variants in the dataset.

We verified this model by using a computer program to generate samples drawn from a randomly mating, constant-sized population generated using the Wright-Fisher model with an infinite-sites model of mutation (Hudson, 2002). Figure 1 shows the expected number of giSNPs as a function of MAF for simulations using six different sample sizes. For all sample sizes, the mean number of giSNPs is in very close agreement with the expectation according to our theoretical model. Figure 1 shows intervals that indicate the range in the number of giSNPs across simulated allelic configurations for each MAF. These intervals are noticeable only for giSNPs occurring at a very low, but non-zero, rate and arise due to the increased sampling variation present for very rare events.

**Figure 1:**
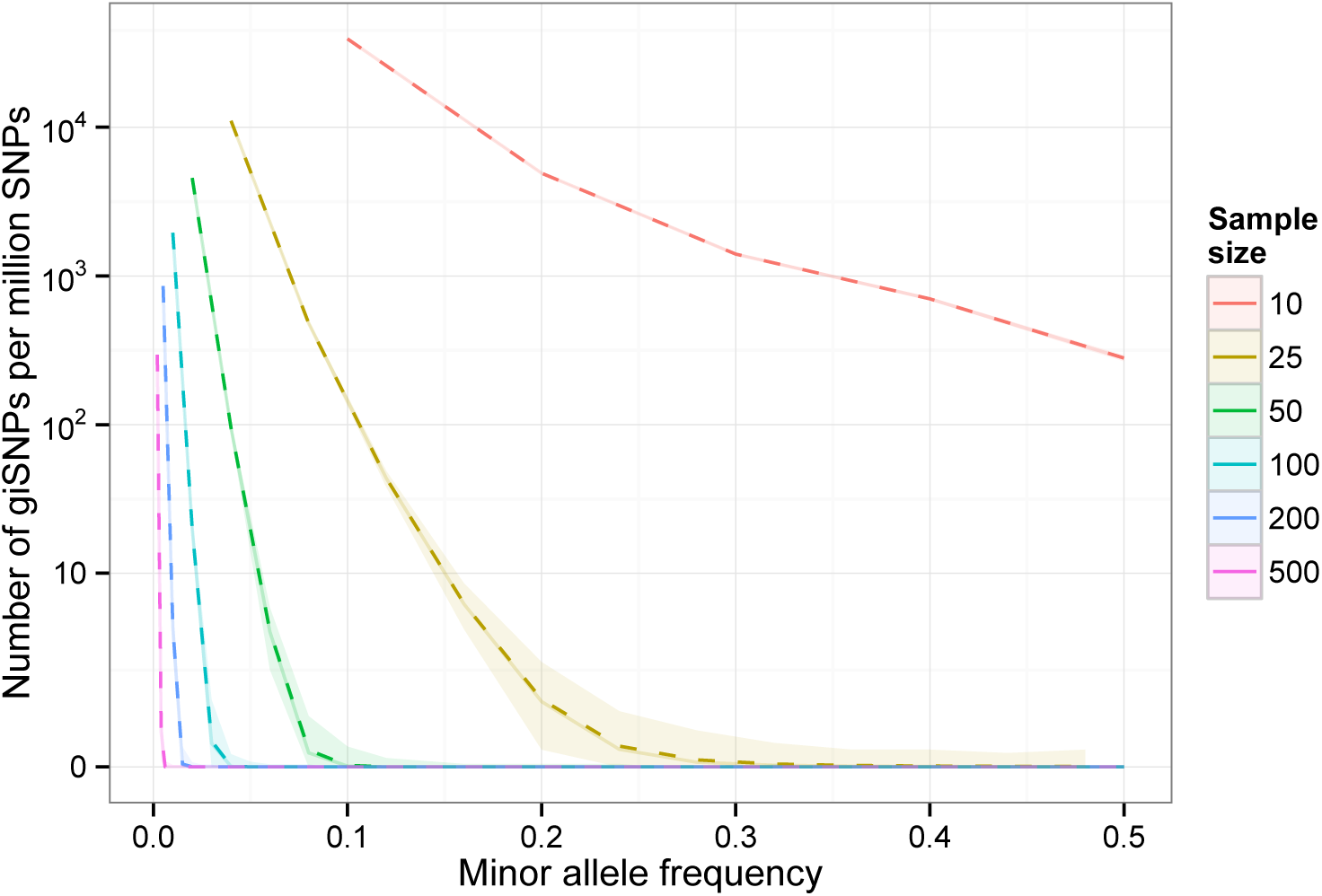
Correspondence between simulated and theoretical expectations of giSNP prevalence. Dotted lines show results calculated using the theoretical model introduced above. Solid lines (directly underneath dotted lines) indicate mean number of giSNPs per million SNPs obtained via simulation (averaged across all sampled allelic configurations). Lighter ribbon surrounding each line shows the range in the number of giSNPs across sampled allelic configurations for each minor allele frequency. Y-axis is inverse hyperbolic sine transformed in this figure and in other figures plotting giSNP prevalence as a function of MAF throughout the paper

In the supporting information, we detail how our theoretical model can be extended to diploid populations. Briefly, in the context of our analysis of giSNPs, we are primarily interested in physically distant pairs of loci (loci that are far apart or on different chromosomes), so phase becomes arbitrary. For unphased allelic configurations that contain no heterozygotes, the expected number of giSNPs is identical to Equation 2, but where N refers not to the number of haploid individuals but to the number of *chromosomes* (*N*/2 diploid individuals). For unphased allelic configurations that contain at least one heterozygote, this probability will be inflated by a factor of 2*^h^*, where *h* is the number of heterozygotes present in the configuration (Supporting Information).

### Key insights from theoretical model

Figure 1 clearly illustrates two central determinants of giSNP prevalence. First, it is clear that the prevalence of giSNPs is reduced as MAF increases. This follows from intuition, as can be shown by considering the extreme example of a singleton SNP present in only one individual: this SNP will have the lowest possible MAF, and will be a member of the giSNP cluster containing all other singleton SNPs present in the same individual. As the MAF increases, the number of possible allelic configurations grows faster than exponentially, and the chance of two unlinked SNPs sharing the same allelic configuration becomes very small for a randomly mating population.

Second, it is apparent from Figure 1 that expectations of giSNP prevalence are strongly dependent on sample size, with almost no giSNPs for sample sizes 200 and 500 except at singleton sites. Again, this phenomenon follows from our theoretical model – the number of possible allelic configurations increases very rapidly as sample size increases, regardless of MAF (although the increase is more dramatic for higher MAF). Overall, these two points indicate that sample size and MAF both play an important role in determining the prevalence of giSNPs through their impact on the total number of possible allelic configurations, a relationship that follows from the combinatorics of minor alleles being partitioned among individuals.

### Demography and mating patterns can exacerbate giSNP prevalence

The theoretical model described above provides quantitative estimates of the prevalence of giSNPs in a randomly mating population of constant size. However, any real population is likely to violate one or more of these assumptions. The derivation above illustrates the strong dependence of giSNP prevalence on sample size and MAF, as discussed previously, but also suggests conditions under which the occurrence of giSNPs is likely to deviate from idealized conditions. We explore these conditions using coalescent simulations with a sample size of *N* = 50. We restrict our simulations to neutrally evolving loci and do not investigate cases of selection or other phenomena such as epistasis. In our simulations, the specific parameters of each departure from the constant-sized panmictic model are necessarily arbitrary, but they will nevertheless be useful for illustrating overall patterns of changes in the prevalence of giSNPs.

### Perturbation of allele frequency spectrum

In the derivation of our model, equation 1 represents a contribution to the prevalence of giSNPs due to the allele frequency spectrum. The allele frequency spectrum is well-known to be affected by both demographic and selective forces (Marth *et al*., 2004; Achaz, 2009). We explored the effect of perturbations to the allele frequency spectrum on giSNP prevalence by simulating data from a population subject to (1) a bottleneck starting a fixed number of generations ago and lasting until the present, and (2) a period of exponential growth starting a fixed number of generations ago and lasting until the present. In the former case the starting population size was constant but the strength of the bottleneck (fraction by which the population size was reduced) varied, while in the latter case the starting population size and magnitude of exponential growth varied such that final population size was constant in all simulations.

As Figures 2A and 2C show, the allele frequency spectrum is distorted by these demographic events. In the case of a strong population bottleneck, a large portion of rare variation is lost and the frequency spectrum shifts toward higher frequency variation (Fig. 2A). When the population grows exponentially at a high rate, the reverse is true and the fraction of rare variation increases dramatically at the expense of common variation (Fig. 2C). Figures 2B and 2D demonstrate that these shifts in the allele frequency spectrum have detectable effects on the prevalence of giSNPs, although they tend to be mild. Specifically, strong exponential growth shifts the allele frequency spectrum toward low frequency variation, which simultaneously creates a larger “pool” of low frequency variants and a smaller pool of higher frequency variants, resulting in slightly higher/lower rates of giSNP occurrence, respectively (Fig. 2C-D). In contrast to exponential growth, a bottleneck has an opposing effect on the allele frequency spectrum in that it produces shifts toward moderate/high frequency variation and away from low frequency variation (Fig. 2A). Perhaps counterintuitively, strong bottlenecks do not lead to any appreciable increases in giSNP prevalence. The reduction in low frequency variants due to the bottleneck decreases giSNP prevalence among low MAF variants (Fig. 2B), but a corresponding increase in higher frequency variants that would be expected to increase giSNP occurrences is mitigated by the vast number of allelic configurations at these MAFs. In particular, the allele frequency spectrum shift toward higher frequency variation (Fig. 2A) does not manifest until MAF ~ 15% (where there are already 100 million possible allelic combinations), which results in a negligible increase (about 1 × 10^−5^ per million SNPs) in the total rate of giSNP occurrence (Fig. 2B).

**Figure 2:**
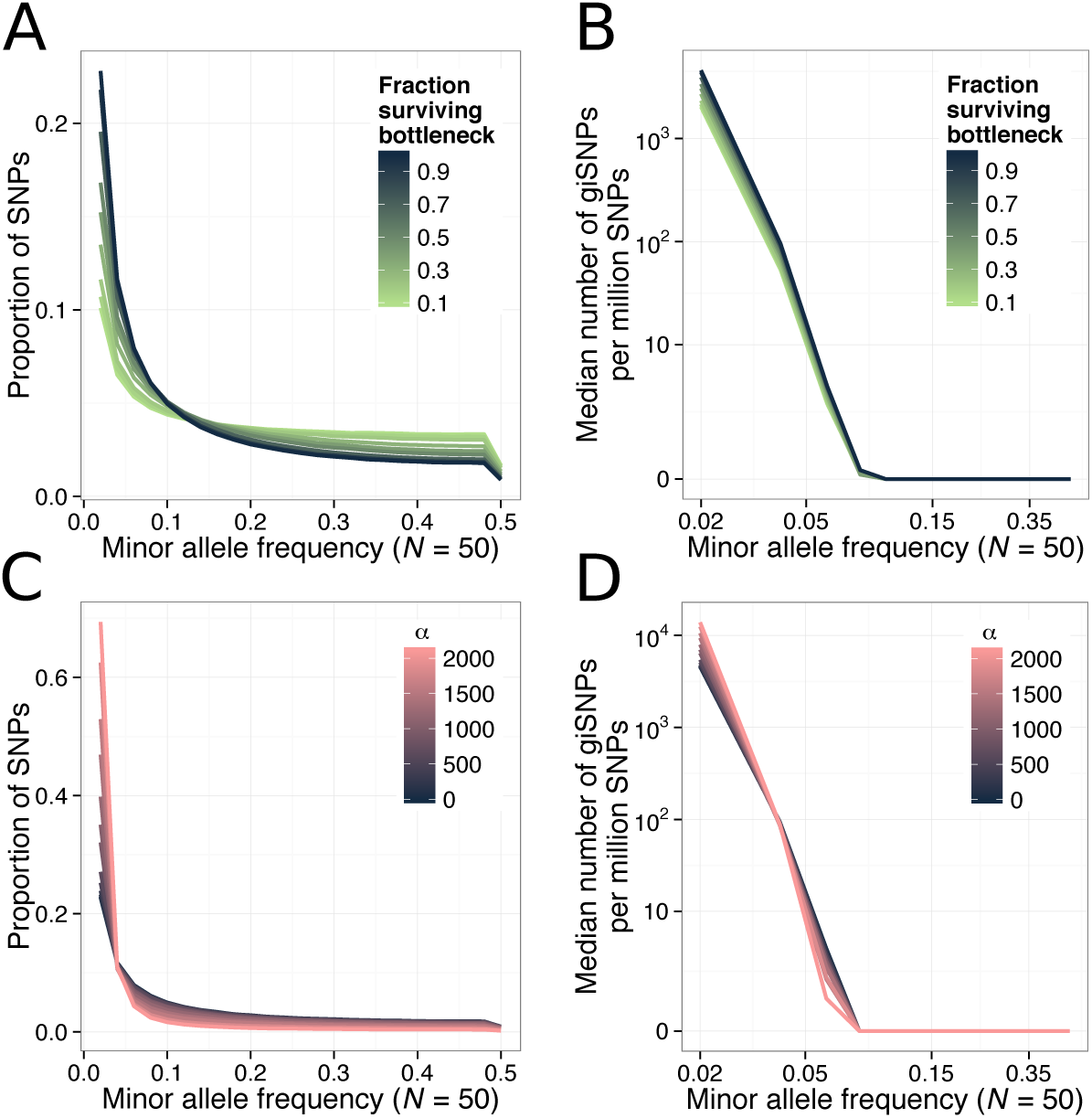
Demographic perturbations can affect the occurrence of giSNPs. Figure shows the allele frequency spectrum and corresponding rate of giSNP occurrence for two demographic models. (A, B) Results from the bottleneck model described in the main text. (C, D) Results from the exponential growth model discussed the main text. In the exponential model, population growth is proportional to exp^-at^ for t units of time; *α* in panels C and D refers to this exponential growth factor.

### Non-uniform probability distribution of allelic configurations

Next, we consider the second component of equation 2, 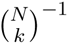. As described above, this factor accounts for the equal likelihood of each allelic configuration, which is a consequence of arbitrary labeling of individuals under the neutral Wright-Fisher model. However, the equal likelihood of all allelic configurations will not hold if mating is non-random. To explore the effect of changes in the probability distribution of allelic configurations due to deviations from random mating among the individuals in a sample, we simulated two models that incorporate population structure: (1) a model of a population that instantly splits into two completely isolated equally-sized subpopulations, and (2) a model of a population that instantly fragments into ten equally-sized subpopulations at a single timepoint with subsequent migration between subpopulations. For the former model we varied the timing of the split, and for the latter model we varied the amount of migration between subpopulations after the fragmentation event. For both models, we drew an equal number of chromosomes from each simulated subpopulation and combined them into one large sample in order to calculate occurrences of giSNPs.

To quantify the non-uniformity of the probability distribution of allelic configurations for a given demographic scenario, we calculated entropy (the “entropy of allelic configurations”) for each MAF. In an information-theoretic sense, the entropy is maximized when the probability distribution is most disordered (a uniform probability distribution of allelic configurations) and minimized when the probability distribution has no randomness (only a single allelic configuration). As Figures 3A and 3C show, the demographic scenarios outlined above produce shifts in the entropy of allelic configurations at only a subset of minor allele frequencies. In the two population model, entropy is drastically reduced for MAF = 0.5, which corresponds to variation either fixing in one subpopulation and being lost in the other or to variation that arose in one subpopulation after the split and subsequently rose to fixation (a “subpopulation-specific” allelic configuration; Fig. 3A). Allelic configurations with MAF slightly less than 0.5 show a reduction in entropy as well, although it is less dramatic. This reduction is due to the overrepresentation of allelic configurations with a small number of differences from the subpopulation-specific configuration. In the ten population model, there are similar dips in entropy that occur at regular intervals rather than only at MAF = 0.5 (Fig. 3C). Specifically, these reductions occur at 10%, 20%, 30%, 40%, and 50% MAF and represent allelic configurations corresponding to variation that is unique to one subpopulation or fixed only within certain subpopulations (with population size 50, 5 chromosomes were sampled from each subpopulation, and 5/50 = 10%, 10/50 = 20%, etc.). Like the two population model, MAFs *near* those corresponding to perfect subpopulation-specific division of variation also display reductions in entropy. As described above, these reductions are due to the overrepresentation of allelic configurations with a small number of differences from the subpopulation-specific configurations. Figures 3B and 3D demonstrate that these shifts in the probability distribution of allelic configurations can have very strong effects on the prevalence of giSNPs. Specifically, a reduction in the entropy of allelic configurations increases the rate of giSNP occurrence for alleles of the same MAF (but does not affect the rate of giSNP occurrence for alleles of other MAFs). Intuitively, if only a few different allelic configurations dominate a particular MAF, the chance of any two variants with that MAF sharing an allelic configuration is much higher than if there were many allelic configurations present at roughly equal frequencies.

**Figure 3:**
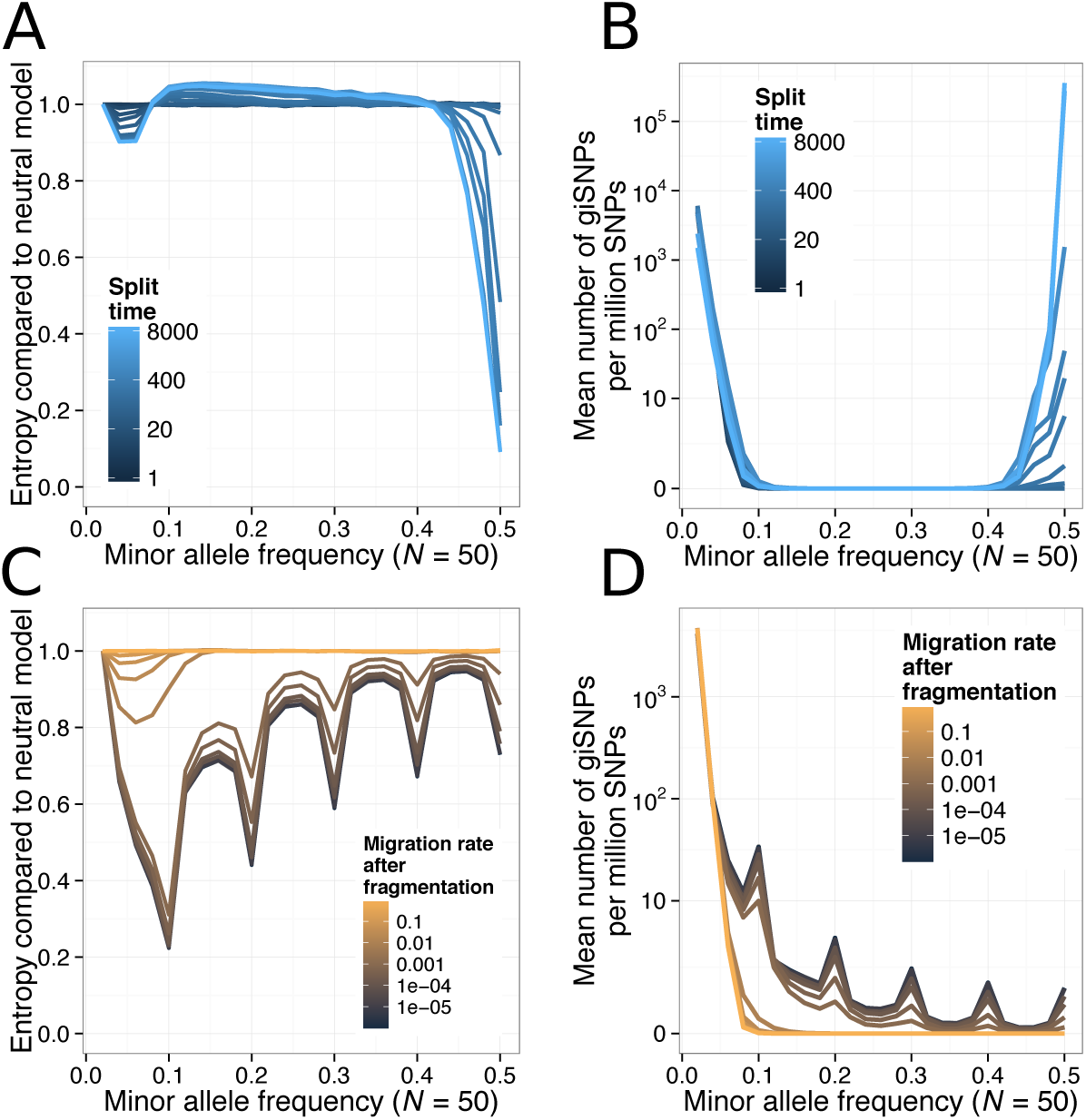
Population structure can greatly increase giSNP prevalence for specific MAFs. Figure shows results from two models that include population structure as described in the main text. (A) and (C) Ratio of the entropy of the distribution of allelic configurations, at each MAF, compared to the neutral scenario (split time of 0 or complete migration between populations, respectively). (B) and (D) Mean number of giSNPs per million SNPs as a function of MAF. Lines in A and B are colored according to split time in generations. Lines in C and D are colored according to the migration rate between populations after the fragmentation event, expressed as the fraction of each subpopulation replaced with migrants each generation.

Overall, these results indicate that population structure can increase the prevalence of giSNPs for particular minor allele frequencies by increasing non-uniformity of the probability distribution of allelic configurations. The particular MAFs that will be affected are specific to the demographic history of the population in question. In the simulated scenarios above, the allele frequency spectrum is also shifted from the neutral case, although this is not strictly required in order to affect giSNP prevalence. We note that there are many other possible scenarios through which deviations from random mating could lead to a non-uniform probability distribution of allelic configurations, including (but not limited to) associative mating, selfing/inbreeding, and other forms of population subdivision. More generally, it is clear from both theory and simulation that non-uniformity of the distribution of allelic configurations, no matter the mechanism, has the potential to be a more important determinant of prevalence of giSNPs than shifts in the allele frequency spectrum. A large shift in the allele frequency spectrum might lead to a change in the fraction of variants at a particular MAF of 2-3 fold. In contrast, strong favoring of a small number of allelic configurations could change the probability of particular allelic configurations by several orders of magnitude.

### An examination of giSNPs in real datasets

We first examined giSNPs in humans using data from the 1000 genomes project (The 1000 Genomes Project Consortium, 2012). We found very few giSNPs, indicating that this phenomenon is unlikely to affect results in modern human GWAS with large sample sizes (Supporting Information). However, we found a very different pattern when we examined the landscape of giSNPs in several large datasets of commonly used model organisms.

Specifically, we comprehensively surveyed LD at all pairwise combinations of SNPs in 100 *Saccharomyces cerevisiae* genomes, 181 *Drosophila melanogaster* genomes, and 206 *Arabidopsis thaliana* genomes (Table 1). We treated all genotypes as haploid, since the yeast genomes sequenced were haploid or homozygous diploid (Strope *et al*., 2015), and the fly and *A. thaliana* individuals were inbred to complete or nearly complete homozygosity (Long *et al*., 2013; Huang *et al*., 2014). In total, our survey of LD at pairwise combinations of SNPs consisted of searching for giSNPs at nearly 12 trillion pairs of SNPs. We found the presence of giSNPs in all datasets, although the amount varied between datasets. We identified billions of giSNPs in the yeast, fly, and *A. thaliana* genomes (Fig. 4A-C, Table 2). Despite this large number of giSNPs, in all three datasets it represented only a tiny fraction (< 1%) of the enormous number of possible pairs of SNPs. In all datasets the bulk of the giSNPs we identified were between variants with low MAF, as predicted by the theory above (Fig. 4A). Nevertheless, we identified thousands of giSNPs with at least moderate allele frequency (≥ 5%; Table 2). According to most measures, the rate of giSNP occurrence was highest in yeast, intermediate in *A. thaliana*, and lowest in flies (Fig. 4A-B).

**Figure 4:**
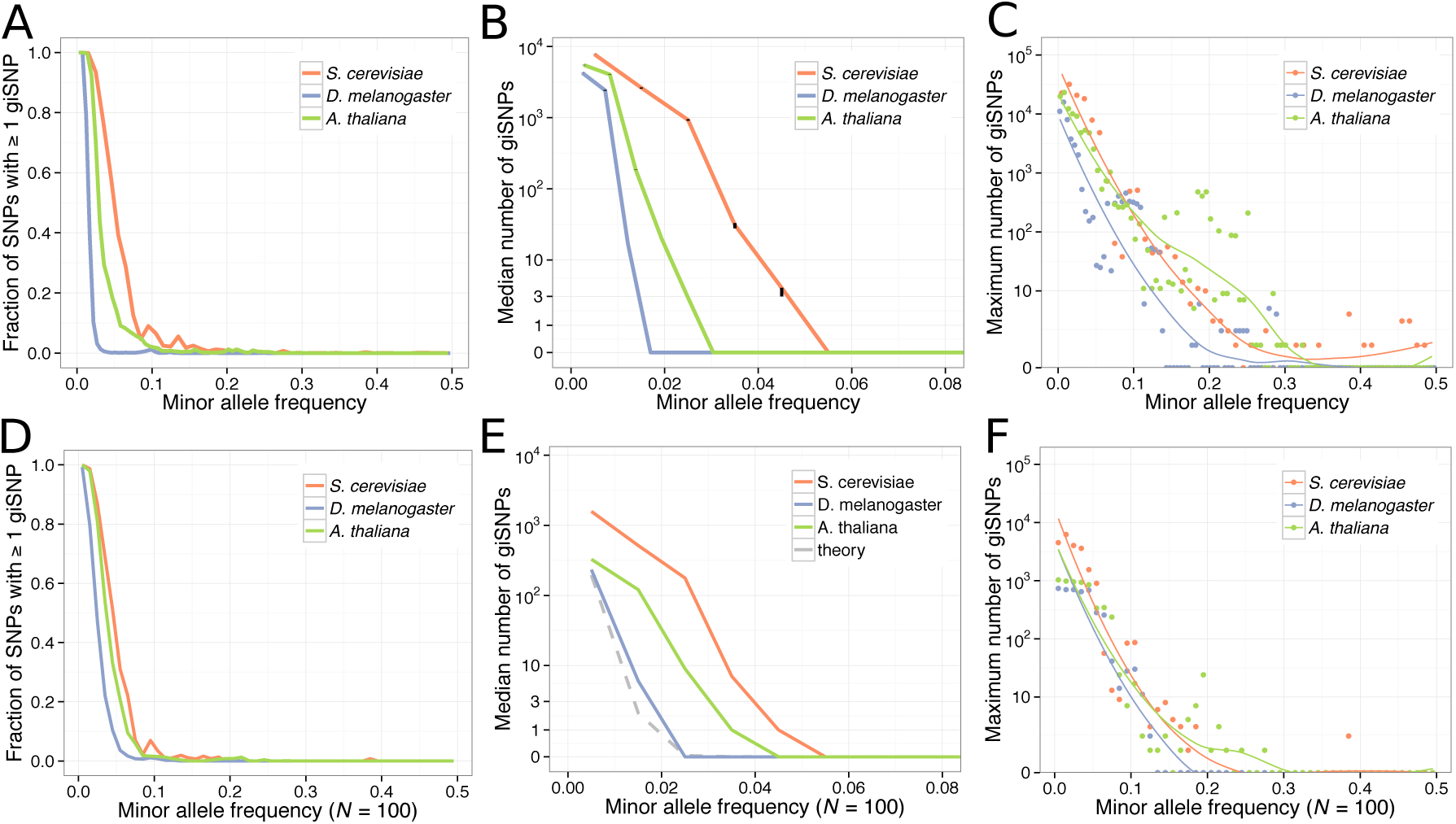
Genetically indistinguishable SNPs in real datasets. (A-C) Statistics calculated from all data for each organism. (D-F) Statistics calculated from datasets downsampled randomly to match a sample size of 100 chromosomes and 100,000 SNPs. (A, D) Fraction of SNPs with at least one giSNP as a function of MAF. (B, E) Median number of giSNPs as a function of MAF. Small black notches indicate bootstrap 95% confidence intervals on the median. The median number of giSNPs for all MAFs > 0.08 is negligible in all datasets. (C, F) Maximum number of giSNPs across all allelic configurations, as a function of MAF. Dots indicate the number of giSNPs for the “worst” allelic configuration at each specific MAF. Solid lines provide a local smoothing via the loess method.

**Table 1:** Characteristics of datasets examined^1^

| Organism | Sample size<sup>1</sup> | Number of SNPs examined | Reference |
| --- | --- | --- | --- |
| *A. thaliana* | 206 | 3,067,372 | Long *et al.* (2013) |
| *D. melanogaster* | 181 | 3,660,570 | Huang *et al.* (2014) |
| *S. cerevisiae* | 100 | 503,846 | Strope *et al.* (2015) |

**Table 2:** Number of giSNPs in real data

| Organism | Number (percent) of possible pairs of SNPs that are genetically indistinguishable |  | Number (percent) of SNPs that have at least one giSNP |  |
| --- | --- | --- | --- | --- |
| | all SNPs | MAF ≥ 5% only | all SNPs | MAF ≥ 5% only |
| *A. thaliana* | $2.72 \times 10^9$ (0.058%) | 38,033 ($< 0.01\%$) | $1.57 \times 10^6$ (51%) | 30,964 (2.5%) |
| *D. melanogaster* | $1.90 \times 10^9$ (0.028%) | 80,936 ($< 0.01\%$) | $1.36 \times 10^6$ (37%) | 1,555 (0.097%) |
| *S. cerevisiae* | $1.22 \times 10^9$ (0.96%) | $5.12 \times 10^6$ (0.052%) | 352,235 (70%) | 10,447 (7.4%) |

A list of pairs of giSNPs could consist of many pairs of independently genetically indistinguishable SNPs, or could be large clusters of a relatively smaller number of SNPs that are all mutually giSNPs. To explore the fraction of SNPs impacted by genetic indistinguishability, we calculated the number of SNPs that have at least one giSNP. We found that a significant fraction of SNPs in each dataset had at least one giSNP, with this quantity especially high in the yeast data (~ 70%; Table 2). The median number of giSNPs per SNP was also highest in yeast, with a slightly lower number in *A. thaliana*, and the lowest number in flies (Fig. 4B). In all cases, the great majority of SNPs that had at least one giSNP were rare (MAF < 5%), although there are thousands of SNPs in each dataset with at least moderate allele frequency that have at least one giSNP (Table 2). To explore the “worst-case” scenario, we tabulated the maximum number of giSNPs among all allelic configurations at each MAF and found in each dataset some configurations with relatively high MAF (> 10%) that had hundreds of mutually genetically indistinguishable SNPs (Fig. 4C).

Overall, our estimates of giSNP prevalence in real datasets are likely to be very slight overestimates due to missing genotype data — we focused on sites with < 10% missing data and ignored individuals with a missing genotype for either SNP in the pair, but it is possible that a missing genotype could break up perfect LD between an otherwise genetically indistinguishable pair of SNPs. We note that, for the *Arabidopsis* data we used, a careful analysis of long-range LD that specifically employed methods to correct for population structure in LD calculations results in many fewer instances of high-LD pairs (Long *et al*., 2013). However, in our examinations of giSNPs in real data, we chose to naively combine individuals known to vary in relatedness in order quantify giSNP prevalence in a set of individuals that might serve as a sample population for GWAS.

### Genetically indistinguishable SNPs in datasets with equal sample sizes

Results from our theoretical model of genetically indistinguishable SNPs and the coalescent simulations described above show that two central determinants of the prevalence of giSNPs are sample size and MAF. In order to examine giSNPs in real datasets on an equal footing from the perspective of sample size, we considered 100 randomly selected individuals from each model organism dataset (*S. cerevisiae, D. melanogaster,* and *A. thaliana*). Since the absolute number of giSNPs is highly dependent on the number of SNPs in a dataset, we randomly selected 100,000 SNPs from each reduced sample of individuals. After removing differences in sample size, the four datasets show similar overall rates of giSNP prevalence (Fig. 4D, F). Nevertheless, there was a consistent pattern of the highest median number of giSNPs being present in yeast and the lowest in flies (Fig. 4E). The theoretical model introduced earlier predicts giSNP prevalence to be roughly on par with that found in flies (Fig. 4E, dashed gray line). This result is consistent with the fact that the *D. melanogaster* dataset is the only dataset we examined that was collected from a single geographic location and could reasonably be thought to derive from a quasi-randomly mating population (Huang *et al*., 2014).

### Almost genetically indistinguishable SNPs

In this paper we have focused on the simplest possible scenario of LD between two independently segregating loci, namely the occurrence of perfect LD between alleles at the two loci. This “worst-case” scenario provides a useful point of entry for studying LD between independent loci, as it is well-suited to simple theoretical modeling and calculations do not depend on the specific measure of LD used. However, LD between alleles at two loci can be substantial without the presence of perfect disequilibrium. We use the term “almost genetically indistinguishable” to describe this phenomenon of high but not complete LD between alleles at two loci, and the abbreviation “almost giSNP” to encompass SNPs that are either genetically indistinguishable or nearly so. Almost genetically indistinguishable SNPs could still present a significant impediment to the localization of causal alleles in the association study design, and might be of more general interest given the relative rarity of perfect LD.

As a preliminary exploration of the scale of almost genetically indistinguishable SNPs relative to giSNPs, we focused on the yeast data described above and quantified LD between all pairs of variants using *r*^2^. As Figure 5 shows, the fraction of SNPs at each MAF with at least one almost giSNP increases as the threshold for disequilibrium decreases from *r*^2^ = 1 (giSNPs) to *r*^2^ = 0.6. Similarly, the mean number (as a function of MAF) and maximum number (across all allelic configurations) of almost giSNPs both increase as the *r*^2^ threshold decreases (data not shown). The maximum MAF at which a substantial fraction of SNPs has at least one almost giSNP is shifted upwards as *r*^2^ decreases; for example, the maximum MAF where at least 25% of SNPs have a giSNP is 6.5% for *r*^2^ = 1, 8.5% for *r*^2^ = 0.8, and 13.5% for *r*^2^ = 0.6 (Figure 5). Interestingly, there is an upward trend in the fraction of SNPs with at least one giSNP that is particularly apparent at the lowest threshold (*r*^2^ = 0.6; Figure 5). This effect likely arises for combinatoric reasons; specifically, assuming that a maximum of *n* alleles can be “flipped” from ancestral to derived or vice versa, there are more possible ways to achieve *r*^2^ = 0.6 starting with an allelic configuration with MAF 0.5 than there are when starting with an allelic configuration of MAF 0.25.

**Figure 5:**
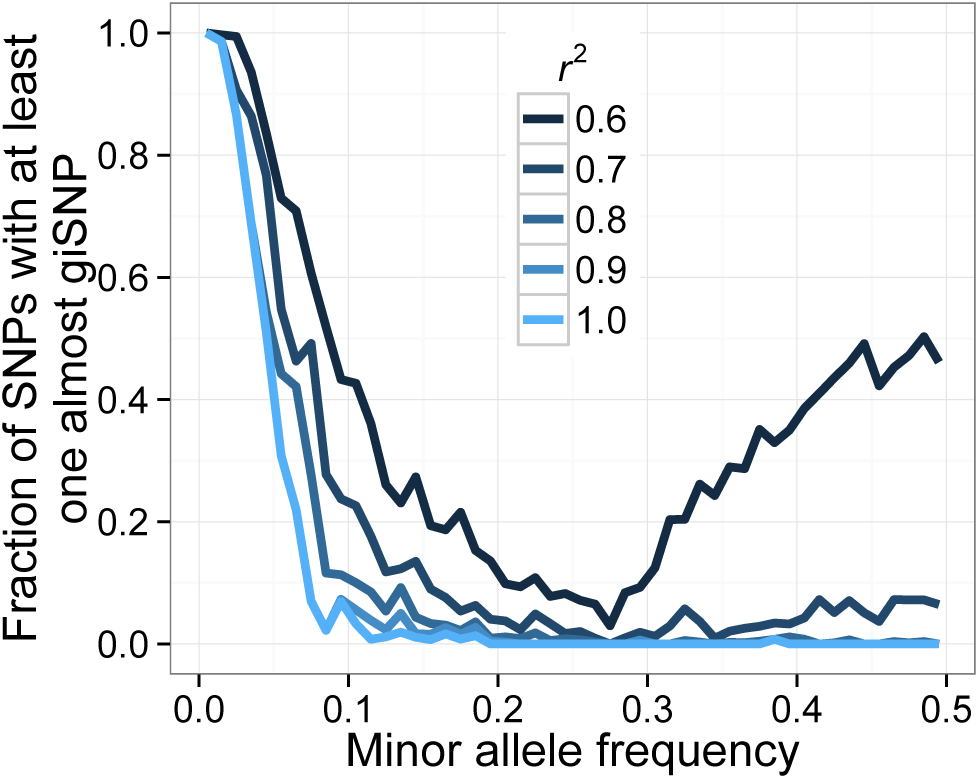
Almost genetically indistinguishable SNPs in 100 *S. cerevisiae* genomes. Lines show the fraction of SNPs with at least one giSNP (*r*^2^ = 1) or almost genetically indistinguishable SNP (*r*^2^ < 1) as a function of MAF.

## Conclusions and Implications for Future Studies

The phenomenon of unlinked genetically indistinguishable SNPs is of practical importance to investigators conducting mapping studies using dense polymorphism data in model and non-model organisms using the association study design. In particular, causal loci with giSNPs elsewhere in the genome could completely eliminate the ability to localize causal alleles via association. Even for Mendelian or nearly Mendelian traits that are commonly considered “easy” to map, giSNPs could lead to confusion about the true source of signal driving a phenotypic association.

As described above, we used theoretical models, simulations, and genome-wide SNP data to explore the conditions under which giSNPs are expected to manifest, and to quantify their prevalence in real data. We find that sample size and MAF are the most important factors determining how often loci are genetically indistinguishable. Nevertheless, giSNPs can also be strongly elevated for particular MAFs under models with deviations from random mating. These observations are particularly relevant for marker-trait association studies in non-model organisms or those where phenotype data is labor-intensive to obtain, as these studies are likely to be conducted with modest sample sizes. In such studies, a global survey of LD could reveal the extent to which it is likely to ultimately impede localization of causal alleles. Such surveys are particularly important for samples consisting of individuals likely to have a complex demographic history, given the potential for strong population structure to elevate the prevalence of giSNPs. At the same time, we recognize that the use of mixed models to account for population structure in association mapping is likely to significantly ameliorate the effect of giSNPs that lie along major axes of population subdivision.

How can we “cure” genetic indistinguishability? In order to reduce the number of giSNPs in a particular dataset, we must break up associations between alleles at two loci that are in perfect disequilibrium. Aside from collecting more individuals (which may not always be possible), the most straightforward way to accomplish this is to take advantage of meiosis, during which recombination and independent assortment break up associations between alleles at loci on the same and different chromosomes, respectively. Specifically, if variation involved in giSNPs is segregating in a cross, and n offspring are collected, then a fraction 1 − 0.5*^n^* of the genetically indistinguishable pairs of loci should be broken up and no longer genetically indistinguishable (assuming giSNPs segregate independently). Although this strategy addresses the “worst-case” scenario of giSNPs, it is not a panacea. As we have shown, almost genetically indistinguishable SNPs can be widespread in genomic datasets. Both genetically indistinguishable and almost genetically indistinguishable SNPs could impede localization of causal alleles. In theory, almost genetically indistinguishable SNPs could result in a stronger marker-trait association between the phenotype and a non-causal locus than at the causal locus itself. Given the potentially disruptive effects of almost genetically indistinguishable SNPs, we suggest that further investigation into its prevalence and practical importance is warranted.

## Acknowledgements

We thank David Swofford for helpful comments regarding simulation methodology. This study was supported in part by NIH grants R01 GM098287 and F32 GM110997.

## Footnotes

1 Including the appropriate reference genome.

